# Neuro-molecular characterization of fish cleaning interactions

**DOI:** 10.1101/2021.06.22.449532

**Authors:** S. Ramírez-Calero, J. R. Paula, E. Otjacques, R. Rosa, T. Ravasi, C. Schunter

**Affiliations:** Swire Institute of Marine Science, Division for Ecology and Biodiversity, School of Biological Sciences, The University of Hong Kong, Pokfulam, Hong Kong SAR; MARE – Marine and Environmental Sciences Centre, Faculdade de Ciências da Universidade de Lisboa, Laboratório Marítimo da Guia, Av. Nossa Senhora do Cabo, 939, 2750-374, Cascais, Portugal; Pacific Biosciences Research Center, Kewalo Marine Laboratory, University of Hawai’i at Manoa, Honolulu, HI, USA; Marine Climate Change Unit, Okinawa Institute of Science and Technology Graduate University, 1919–1 Tancha, Onna-son, Okinawa 904–0495, Japan; Australian Research Council Centre of Excellence for Coral Reef Studies, James Cook University, Townsville, Queensland, 4811, Australia

**Author notes:** Correspondence to: Celia Schunter.

**Keywords:** Social Behaviour, Molecular Pathways, Transcriptomics, Species Interaction, Learning and Memory

## Abstract

Coral reef fish exhibit a large variety of behaviours crucial for fitness and survival. The cleaner wrasse *Labroides dimidiatus* displays cognitive abilities during interspecific interactions by providing services of ectoparasite cleaning, thus serving as a good example to understand the processes of complex social behaviour. However, little is known about the molecular underpinnings of cooperative behaviour between *L. dimidiatus* and a potential client fish (*Acanthurus leucosternon*). Therefore, we investigated the molecular mechanisms in three regions of the brain (fore-, mid-, and hindbrain) during the interaction of these fishes. Here we show, using transcriptomics, that most of the transcriptional response in both species was regulated in the hindbrain and forebrain regions and that the interacting behaviour responses of *L. dimidiatus* involved immediate early gene alteration, dopaminergic and glutamatergic pathways, the expression of neurohormones (such as isotocin) and steroids (*e*.*g*. progesterone and estrogen). In contrast, in the client, fewer molecular alterations were found, mostly involving pituitary hormone responses. The particular pathways found suggested learning and memory processes in the cleaner wrasse, while the client indicated stress relief.

## INTRODUCTION

Social behaviour allows species to establish biological relations through intra- and interspecific interactions. These relationships prompt species to generate social mechanisms to survive (*e*.*g*. detect predators), reproduce (*e*.*g*. courtship) and thrive in nature (*e*.*g*. territoriality, living in groups; [1]). Indeed, social behaviour is an ability that promotes responses to specific situations (*i*.*e*. competition for shelter or food), including biotic factors and their physical environment [2,3]. This ability to respond to social stimuli can be regulated to optimize their relationships with conspecifics and other species, allowing them to perform more effectively in nature [1]. At present, the study of social behaviour and its mechanisms have been centred on understanding the capacity to regulate and change social relationships (social plasticity) that can enhance and promote survival [4,5].

Studies on this behaviour have focused on the genetic, epigenetic, endocrine and neural mechanisms underlying social behavioural responses [1,6,7]. For example, studies on the evolution of social phenotypes and transcriptomic signatures in mice, sticklebacks and honey bees, have elucidated on the mechanisms regulated during the response to social challenges (*e*.*g*. territory intrusion) [8,9]. One well studied group of genes are referred to as immediate early genes (IEGs) and are used to detect early neural activation as indicators of adaptive plasticity and learning processes [55,56]. Other groups of genes suitable for understanding social responses (*e*.*g*. cooperation and aggression) are genes regulated in molecular pathways related with the neuron system (*e*.*g*. Dopaminergic pathway), or with the transduction of signals (stimuli) in the brain (*e*.*g*. MAPK signalling pathway) [10–12]. These pathways are modulated during the interactions or communications between individuals, and social behavioural phenotypes can therefore be linked to particular gene expression patterns [11,13]. While many studies focus on major social challenges (*e*.*g*. territory defense, cooperation, dominance), little is known about neuro-molecular responses of other key social interactions such as marine cleaning mutualisms. Cleaning mutualism is one of the most important interactions between coral reef fishes as it involves the removal of ectoparasites of the skin of cooperative hosts and establishing long-term social interactions that are key to maintaining the biodiversity of the ecosystem [14–16]. Therefore, to further understand the gene regulation of this type of social interaction, organisms displaying well-developed social systems and sophisticated cognitive abilities are essential to investigate the functional basis of cleaning mutualisms [1,4,6,17].

The coral reef bluestreak cleaner wrasse *Labroides dimidiatus* is one of the most remarkable examples of mutualistic cleaning behaviour in marine species. It is widely known for enhancing fish biodiversity in local communities due to its important role in cleaning ectoparasites from the skin of hosts and for its complex social behaviour [18,19]. Studies on this species as a model for behaviour, have focused on neural physiological responses in its interaction with other species and how different social situations such as the establishment of social bonds modulate their behaviour [14,20–22]. For example, dopamine and serotonin levels influence the motivation of cleaners to engage in interactions [23] and are induced during social stress [38]. These monoaminergic hormones (dopamine and serotonin) also play a role in the regulation of the service of cleaning in this cleaner wrasse and their willingness to interact, thus modifying the capacity of individuals to react to new social scenarios (*i*.*e*. presence of new clients [5,25]). However, even though these studies have been conducted at behavioural and neurobiological scales, none have evaluated the molecular signatures or gene expression regulation that underlie this type of behaviour.

In this study, we identified the molecular signatures of the interaction behaviour of *L. dimidiatus* with its client fish the powder-blue surgeonfish *Acanthurus leucosternon*, across three major regions of the brain: forebrain (FB), midbrain (MB) and hindbrain (HB). Whole-genome differential gene expression patterns were analysed by comparing both cleaner and client individuals after social cleaning interactions against individuals without interaction (control). We explore the gene expression altered during the interaction and investigate molecular signatures associated with social behaviour. Finally, identifying underlying functional processes altered in these two species contributes to the understanding of mutualistic cleaning behaviour at lower levels of biological organization and exhibit possible mechanisms of social plasticity during interspecific cleaning interactions.

## MATERIALS AND METHODS

### (a) Experimental setup

To test for the molecular mechanisms involved in the interaction between two fish species, 12 female adult individuals of *L. dimidiatus* and 12 female adults of *A. leucosternon* were collected from the wild in the Maldive Islands and transported by TMC-Iberia to the aquatic facilities of Laboratório Marítimo da Guia in Cascais, Portugal (figure 1, table S1a). We selected a common Acanthurid species as client since they are one of the most frequent hosts for *Labroides* species in coral reefs [20]. We also selected female individuals to be consistent with past studies using *L. dimidiatus* as a model species and because gene functions may differ between sexes and can blur the analysis of molecular signals [26]. Fishes were habituated for 28 days to laboratory conditions. Water parameters were monitored daily and automatically controlled using an aquarium computer (Profilux 3.1N GHL, Germany). Seawater conditions were kept similar to their capture site (salinity = 35 ± 0.5, temperature 29 ± 1°C, pH 8.10 ± 0.05). Each cleaner fish was kept separately in individual tanks (20L) to avoid aggression as they are highly territorial animals. In contrast, surgeonfish (*A. leucosternon*) were held in groups of three individuals in 20 L tanks. Fish were fed *ad libitum* once per day. Each experimental tank had a flow-through aquatic system in which levels of alkalinity, dissolved carbon and pH were strictly maintained. Natural seawater was pumped from the sea to a storage tank of 5m^3^ and then filtered and UV-irradiated with a Vecton V2 300 Sterilizer before reaching the experimental tanks. Experimental tanks were kept under a photoperiod of 12 h/12 h (light/dark cycle). Ammonia and nitrate levels were checked daily using colorimetric tests and always kept below detectable levels (Salifert Profi Test, Netherlands). Seawater temperature was regulated using chillers (Frimar, Fernando Ribeiro Lda, Portugal) and underwater heaters 300 W, (TMC-Iberia, Portugal). Salinity was daily monitored with a V2 refractometer (TMC-Iberia, Portugal), and pH and temperature with a VWR pH 1100H pH meter (Avantor, US).

**Figure 1.**
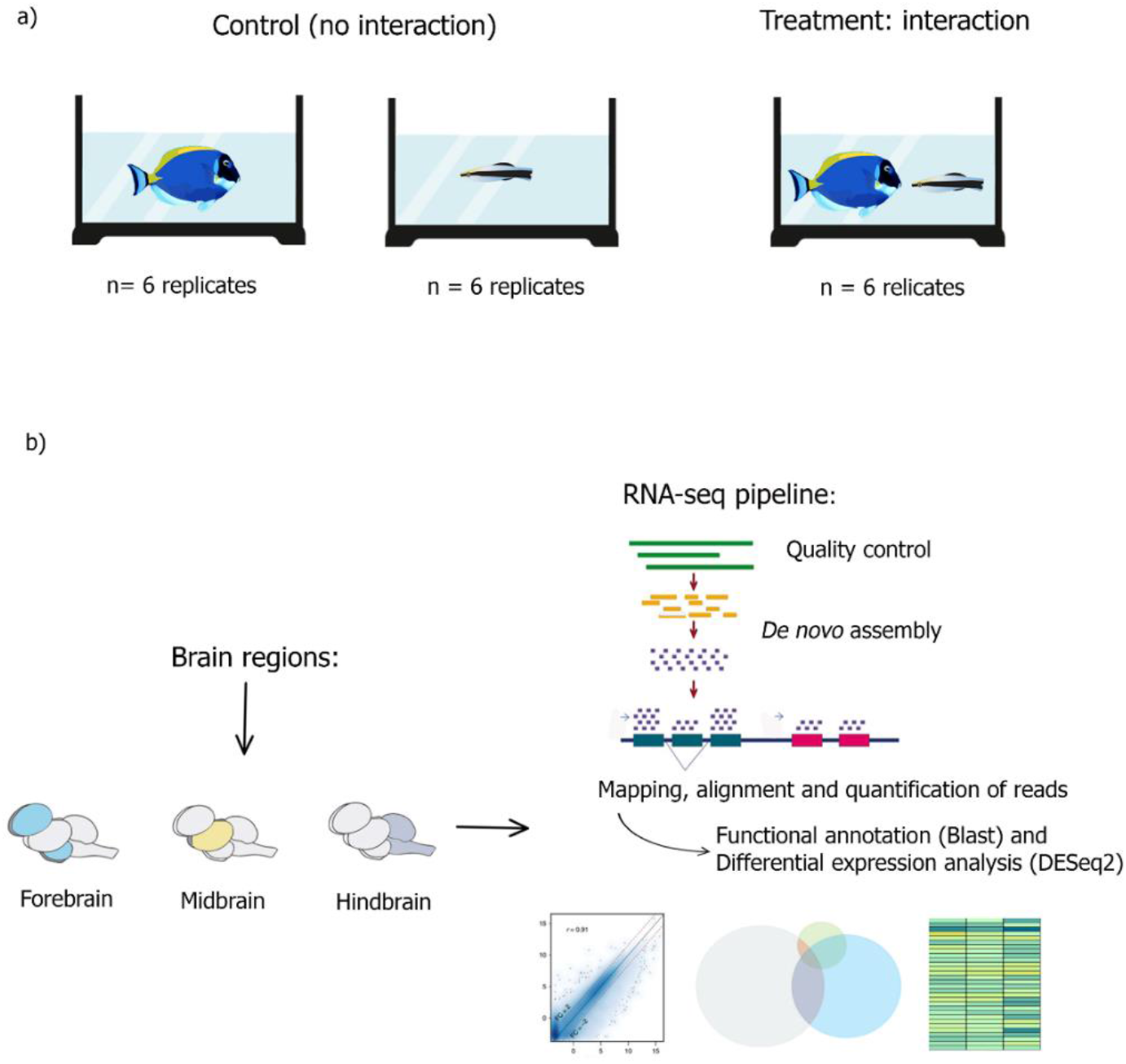
a) Experimental design in which *Labroides dimidiatus* (N=12) and *Acanthurus leucosternon* (N=12) were kept separately (control: no interaction) or allowed to interact (condition: interaction) in the observation tanks (DGAV – Permit 2018-05-23-010275). 40 min of video were recorded. b) Brain regions dissections (forebrain, midbrain and hindbrain) for each species, and RNA-seq pipeline including *de novo* transcriptome assembly, differential gene expression and functional analysis. Further details of the individuals used can found at table S1a.

Behavioural tests started after the 28 days of laboratory acclimation. The tests were performed in observation tanks (40 L) set in an isolated observation room. The fish were either placed into a tank with i) no-interaction (control) or ii) interaction (cleaner-client pair). In the no-interaction treatment, cleaners or clients were kept alone in the observation tank (control), while for the interaction treatment, pairs composed of one cleaner and one client were kept together in the observation tank, allowing them to have close contact (figure 1). Their interactions were filmed for 40 minutes. At the end of the observation period, each fish was immediately euthanized by severing the spine, body length was measured, and three separated regions of the brain were immediately dissected out (figure 1b): forebrain (diencephalon and secondary prosencephalon), midbrain (tectum opticum, torus semicircularis and tegmentum), and hindbrain (cerebellum and medulla oblongata), as behaviour is regulated differently in these regions of the brain in teleost fishes [22]. Finally, tissues were placed in a tube, snap-frozen and kept at -80° C for further processing.

### (b) Behavioural Video Analyses

Behavioural analysis was performed in both treatments. For no-interaction individuals (control), we looked for abnormal behaviour and stress such as erratic movement, secession of swimming or aggressive postures. No abnormal or stress like events were found. For the interaction treatment individuals, we analysed cleaning behaviour according to [27]. Cleaning behaviour was grouped into two categories, i) cleaning interactions & motivation and, ii) interaction quality [23]. To characterise cleaning interactions & motivation, we measured the number of interactions (*i*.*e*. close body inspection and removal of damaged tissue or scales including inspection allowing posture of the client), the number of interactions initiated by both cleaners and client, as well as the number of posing displays the client conducted to attract the cleaner. Interaction quality was determined by the duration of interactions, the number of client jolts (conspicuous signals that indicate cheating or dishonesty by the cleaner [28]), the number of times clients were chased by the cleaners to initiate interactions, and the number and duration of tactile stimulation events (touches with pectoral fins that reduce stress levels and prolong interaction duration [16,29]). All behavioural videos were analysed using the event-logging software “Boris” [30], and further information can be found on table S1b.

### (c) RNA extraction and transcriptome assembly

For RNA extractions 300 ***µ***l of RTL Buffer was added to the frozen tissue with several sterile silicon beads for tissue homogenization in a Tissuelyzer (Qiagen) for 30 seconds at maximum speed to then follow the RNAeasy Mini Kit protocol including a DNase I treatment (Qiagen). The resulting RNA was tested for quality on an Agilent Bioanalyzer and all samples met quality standard of RNA Integrity Number (RIN) > 8. mRNA-focused sequencing libraries were designed with Illumina TruSeq v3 kits and sequenced for 150 bp paired-end reads on an Ilumina Hiseq4000 at the King Abdullah University of Science and Technology corelab facility.

In order to assess the molecular basis to species interactions, on average 31.4 million raw reads for *L. dimidiatus* and 33.8 million for *A. leucosternon* were processed following a bioinformatic pipeline (table S2a-b). Quality was examined using FastQC v. 0.11.9 [31], reads were trimmed and adapters were removed to avoid the presence of poor-quality sequences in our *de novo* transcriptome assembly. For this, we used Trimmomatic v.0.36 [32] with parameters as follows: ILLUMINACLIP:TruSeq3-PE.fa:2:30:10 LEADING:4 TRAILING:3 SLIDINGWINDOW:4:15 MINLEN:40. For both species, there is currently no genome reference available, hence a *de novo* transcriptome assembly was constructed for both species separately after several tests with different number of samples using the default parameters in Trinity v. 2.8.5 [33]. A total of five individuals per species was chosen including the three regions of the brain (table S2a). To assess the quality of the resultant assemblies for each species, we investigated the read representation against our *de novo* assemblies using the aligner Bowtie2 v. 2.3.4.1 [34], with the *–very-sensitive* parameter. To reduce the transcript redundancy and to obtained only coding transcripts, we used the software transdecoder v. 5.5.0 [33] to identify the candidate coding regions ORF (open reading frame), keeping the option -*single_best_only*. From these results, we retrieved the output file with the final candidate ORF regions of more than 100 bp and then conducted a BLAST analysis using the Swissprot/Uniprot and Zebrafish databases obtained from www.uniprot.org (Swiss-Prot: November 2019) and NCBI (*Danio rerio*, txid7955, Apr 2018), respectively. Moreover, we explored the completeness of our assemblies using BUSCO v3 [35] to obtain the number of conserved ortholog content from our results represented in the dataset *Actinopterygiiodb9*. Finally, we computed the Nx statistics to estimate the approximate length of the transcripts in each assembly (N50) using the trinity script *trinity*.*stats*.*pl*. The N50 statistic provides information on the length of at least half (50%) of the total assembled transcripts. Consequently, the assembly for each species with the most complete and longest transcripts and with at least 95% of protein recovery was chosen as a reference (table S3).

We annotated the transcripts of the final *de novo* transcriptome assemblies for each species using BLAST+ 2.10.0: December 16, 2019, and the databases Swissprot/Uniprot protein database (November 29, 2019), Zebrafish (*Danio rerio*, release Apr 2018) for both species, and the Ballan wrasse (*Labrus bergylta*, release March 2020) for *L. dimidiatus* only, obtained from Ensembl (GCA_900080235.1). We used *L. bergylta* because it is the closest species to *L. dimidiatus* with an available genome annotation. Finally, we used Omicsbox v. 1.3 [36] to functionally annotate the transcripts with Gene Ontology (GO terms) and KEGG pathways.

### (d) Differential Expression Analyses

To perform differential expression analyses, we quantified transcript abundance for each species using the trinity script *align_and_estimate_abundance*.*pl* using RSEM v1.3.3 as quantification method and Bowtie2 [34] as mapping tool. Of the final gene expression matrix we filtered out transcripts with no expression by using the script *filter_low_expr_transcripts*.*pl* while retaining the most highly expressed isoforms for each gene using the command *–highest_iso_only*. To statistically evaluate differential gene expression, we used DESeq2-package v. 1.26.0 [37] with a Wald test statistic with a *design = ∼treatment* for each region of the brain separately, an FDR p-adjusted significance value of 0.05 and an absolute log2fold change threshold of 0.3 as a cut-off as previously used in other study evaluating fish brain transcriptomics [38] and to remove further potentially spurious significant differential expression. We compared the individuals from control vs interaction to determine their significant differential gene expression for the three regions of the brain forebrain (FB), midbrain (MB) and hindbrain (HB) to retrieve the molecular response specific to these areas for both species, but separately for each species. Once statistically significant differentially expressed genes (DEGs) were obtained, functional enrichments were performed using Fisher’s exact test by testing the DEG subsets against the whole *de novo* transcriptome with a cut-off of FDR 0.05 in Omicsbox v. 1.3. Each fish species was analysed separately to capture the full breadth of differential gene expression per species due to the fact that the molecular reactions vary across organisms. The GO term IDs obtained were used as a reference to interpret the over-represented or under-represented molecular functions underlying the differentially expressed genes during the interaction, according to the annotations from the universal PANTHER (Protein Analysis Through Evolutionary Relationships) Classification System.

## ETHICAL NOTE

This work was conducted under the approval of Faculdade de Ciências da Universidade de Lisboa animal welfare body (ORBEA – Statement 01/2017) and Direção-Geral de Alimentação e Veterinária (DGAV – Permit 2018-05-23-010275) following the requirements imposed by the Directive 2010/63/EU of the European Parliament and of the Council of 22 September 2010 on the protection of animals used for scientific purposes. A Material Transfer Agreement of biological samples (fish brains) was signed between MARE and KAUST.

## RESULTS

### Behavioural analysis

In the interaction condition every pair (6 replicates) of *L. dimidiatus* and *A. leucosternon* engaged in cleaning interactions. On average, each pair engaged in 43 ± 17.5 interactions, with an average duration of 13 ± 3.6 seconds, corresponding to a proportion of time spent interacting of 13 ± 6.6 % out of the 40 minutes of behavioural trials. Of these interactions, on average, 75 ± 13.7 % were initiated by the cleaner. The cleaner fish’ dishonesty was answered with an average of 3 ± 2.8 client jolts and 2 ± 1.9 chases. Lastly, cleaners used tactile stimulation for reconciliation on average 3 ± 3.4 times. In the control, both *L. dimidiatus* and *A. leucosternon* were swimming around the tank without any display of abnormal behaviour or stress. All behavioural data can be found in table S1b.

### De novo transcriptome assembly

The obtained final de novo transcriptome assemblies are the first references for both *L. dimidiatus* and *A. leucosternon* and resulted in 114,687 and 123,839 coding transcripts respectively (NCBI accession: GJED00000000 & GJGS00000000). Values of N50 indicated that at least half of the transcripts had a length of 1,813 and 2,827 bases, respectively. For *L. dimidiatus* and *A. leucosternon*, a total of 26,380 and 30,770 transcripts were annotated using the Swissprot database, respectively. In addition, a total of 8,379 and 6,438 transcripts were annotated using the *Danio rerio* reference, and 28,988 transcripts were annotated only for *L. dimidiatus* using *Labrus bergylta* database. Further metrics and assembly steps can be found in tables S2 and S3.

### Gene expression analyses

#### Labroides dimidiatus

We evaluated the gene expression differences between individuals from control, which had no interaction, against individuals from the interaction condition. 46 commonly differentially expressed genes (DEGs) exhibiting functions such as dendrite cytoplasm, mRNA binding, sodium ion transmembrane, transporter activity, ATP binding, positive regulation of dendritic spine morphogenesis (figure 2c; figure S1; table S4). When analysing differential gene expression for each of the three brain regions separately, we found that the hindbrain (HB) exhibited the highest number of differential gene expression among the brain regions (2,728 DEGs, figure 2a), followed by the forebrain (FB; 1,414 DEGs) and the midbrain (MB; 421 DEGs). In the HB, 1,370 significantly enriched functions were found, including negative regulation of neuron apoptotic process, synaptic cleft, AMPA glutamate receptor activity, long-term memory, sensory perception of sound, and a total of 14 enriched functions related to behaviour such as motor behaviour, grooming behaviour, behavioural fear response (table S7 and figure S3). In the FB, 980 biological processes were related to behaviour such as signal transduction, calmodulin binding, social behaviour, locomotory exploration behaviour, vocalization behaviour, among others (table S5and figure S1). Finally, for the MB, functional enrichment only resulted in 357 significant functions, including nuclear-transcribed mRNA catabolic process, nonsense-mediated decay, neuronal action potential and sodium ion transmembrane transport (table S6and figure S2). Biological functions regarding behaviour in this brain region were related with behavioural response to pain, locomotory behaviour, male courtship behaviour, maternal behaviour and behavioural defence response (table S6).

**Figure 2.**
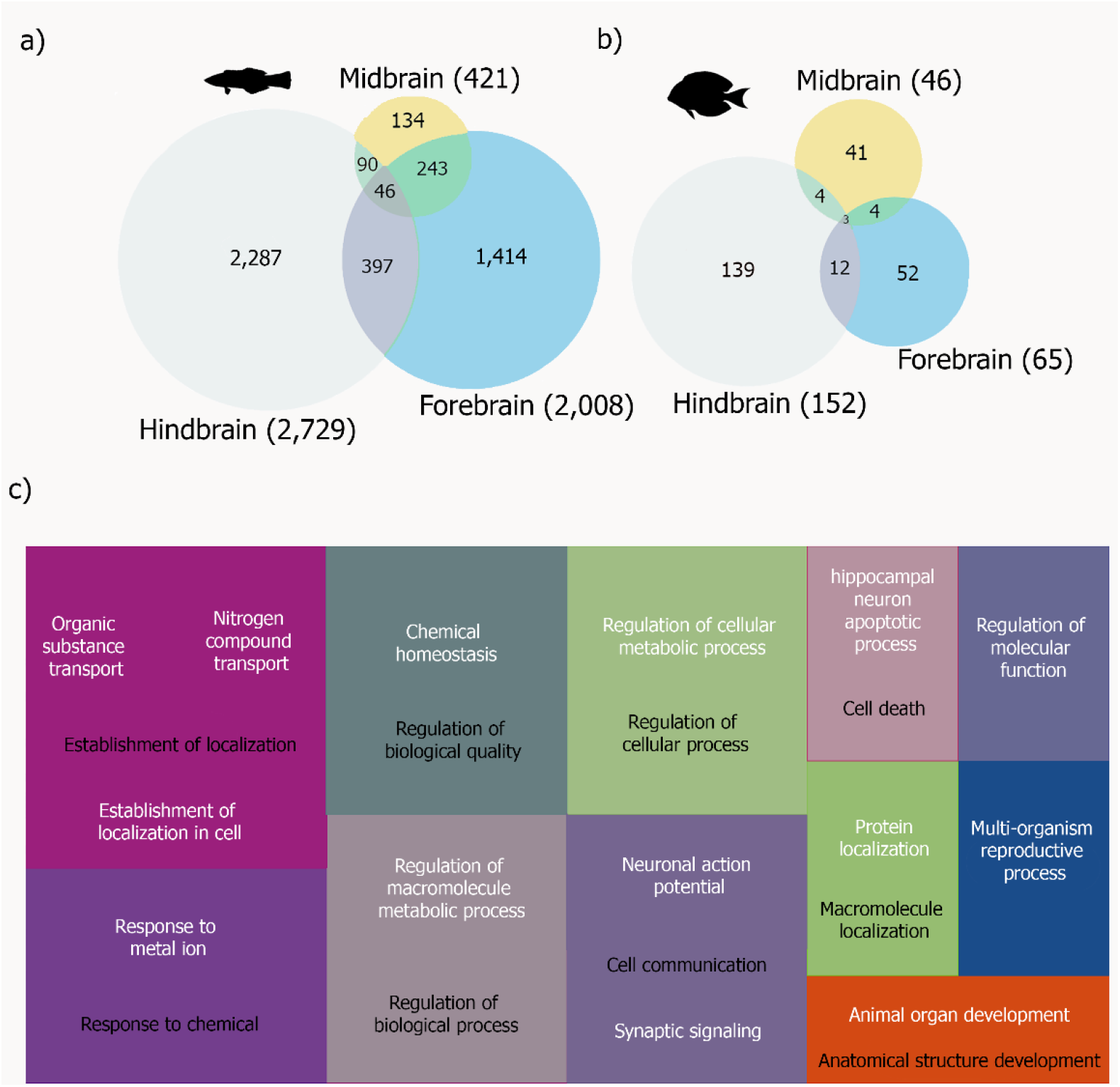
Number of differentially expressed genes between control and interaction for each of the three brain regions and the common overlap of (a) the cleaner fish *Labroides dimidiatus* and (b) the powder-blue surgeonfish *Acanthurus leucosternon*. Numbers in brackets represent the total differential expressed genes found at each brain region. (c) Gene Ontology treemap for *L. dimidiatus* representing the commonly enriched functions across the three brain regions when interacting with a client. Boxes with the same colour correspond to the upper-hierarchy GO-term, and its title is found in the middle of each box. For *A. leucosternon* no common enriched functions were found across the three brain regions. Description of the enriched functions for both species can be found in tables S5-7 and S8-10.

#### Acanthurus leucosternon

For *A. leucosternon*, the total number of DEGs was lower in comparison with *L. dimidiatus*. Only three common differentially expressed genes were found across the three brain regions, which are RNA/RNP complex-1-interacting phosphatase (DUS11), ATP-dependent DNA helicase PIF1 (PIF1) and a third gene for which no annotation was obtained. However, when analysing differential gene expression in each brain region separately, we found 139 DEGs in the HB and nine enriched in functions such as DNA metabolic process, DNA repair, biological regulation and protein binding (table S10). For the FB, 52 DEGs were detected with six enriched molecular functions in signalling receptor activator activity, receptor regulator activity, hormone activity, and two biological functions: gas and oxygen transport (table S8). Finally, 41 DEGs were found in the MB, and only Rab protein signal transduction was significantly enriched (table S9). Like *L. dimidiatus*, the HB exhibited the highest number of DEGs in the comparison of control vs interaction (figure 2a and b).

Evidence of significant differential expression of Immediate Early Genes (IEG) was found in *L. dimidiatus* with 31 differentially expressed IEGs in the three brain regions (figure 3a). However, more differentially expressed genes were found in the HB region (20) and the FB region (16) (figure 3a). Many of the functions of these genes are involved in signalling (*i*.*e*. RHEB, RGS6/9 and 19), transcription factor activation (*i*.*e*. KLF5/11) and plasticity (*i*.*e*. NPAS4L). Differential expression of genes associated with learning processes, memory and neural development was also observed. For instance, genes FOS and SBK1 in the FB and HB, being SBK1 downregulated in both regions, while FOS in the HB only. Moreover, genes KLF11 and JUN were differentially expressed in the HB region (figure 3a; table S11). Finally, for *A. leucosternon*, the only IEG differentially expressed found in our dataset was JUN in the HB, hence found differentially expressed in both studied species.

**Figure 3.**
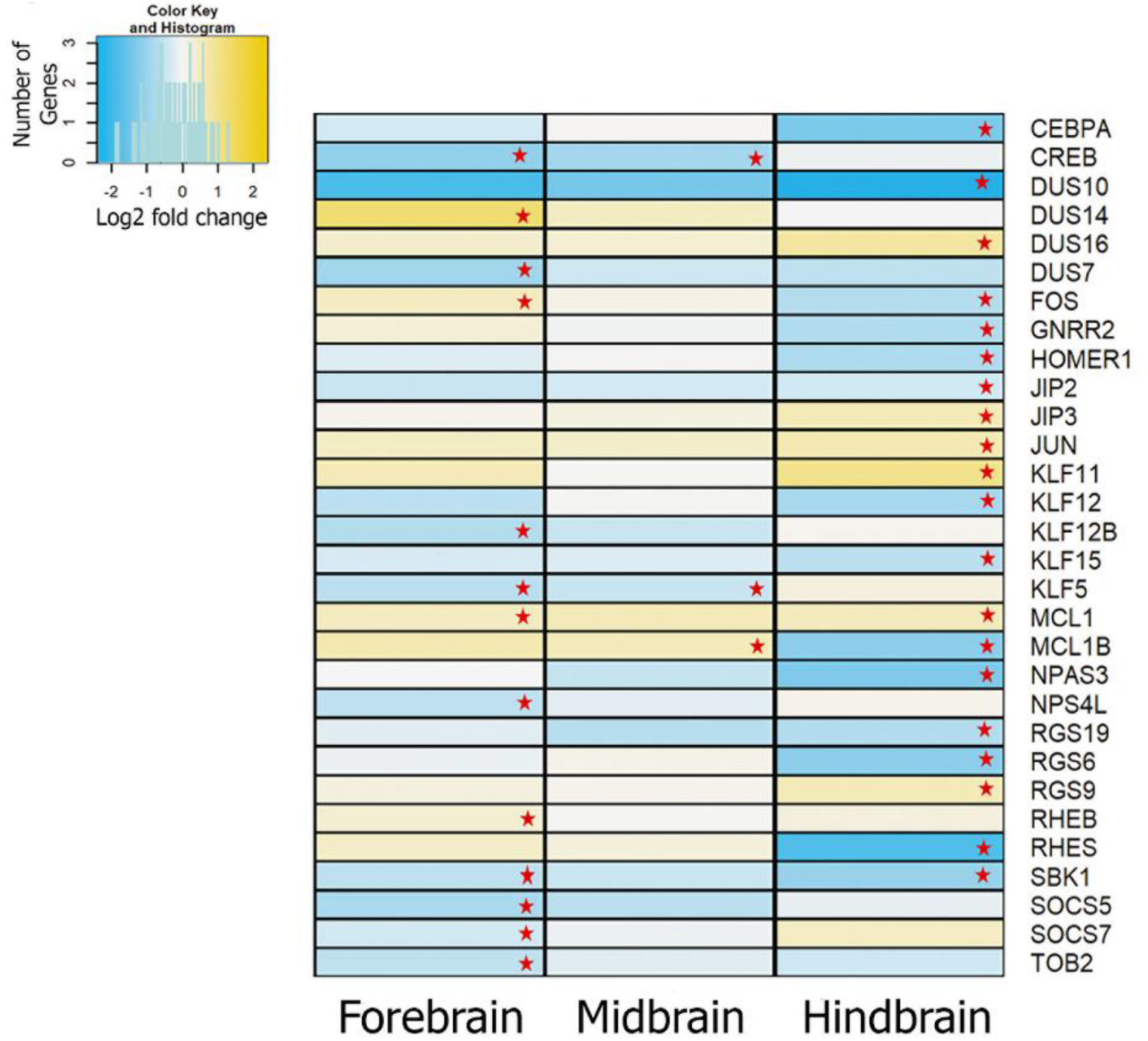
Comparative differential gene expression patterns of Immediate Early Genes between the regions of the brain of *L. dimidiatus*. Red stars represent significance. The legend indicates the reference values of log2fold changes for each DEG in the figure.

Additional groups of genes differentially expressed during the interaction were found. For instance, Dopamine pathway genes such as Dopamine receptors DRD1, 2, 5, Tyrosine 3-monooxygenase (TH), Dopa Decarboxylase (DDC) and Dopamine beta-hydroxylase (DOPO) were differentially expressed in the HB region of *L. dimidiatus* (figure 4a; table S12). These genes were downregulated during the interaction condition, and enriched functions such as the regulation of dopamine secretion and D2 dopamine receptor binding were shared between the FB and HB. Furthermore, Dopamine biosynthesis processes, Dopaminergic pathway, adenylate cyclase-activating dopamine receptor signalling pathway and the regulation of dopamine secretion were specific for the HB, while the regulation of synaptic transmission, dopamine metabolic process, D1 dopamine receptor binding, and the cellular response to dopamine were enriched functions only in the FB for *L. dimidiatus* (figure 4b, figure S4). Finally, no differential expression of these genes was found in the MB region.

**Figure 4.**
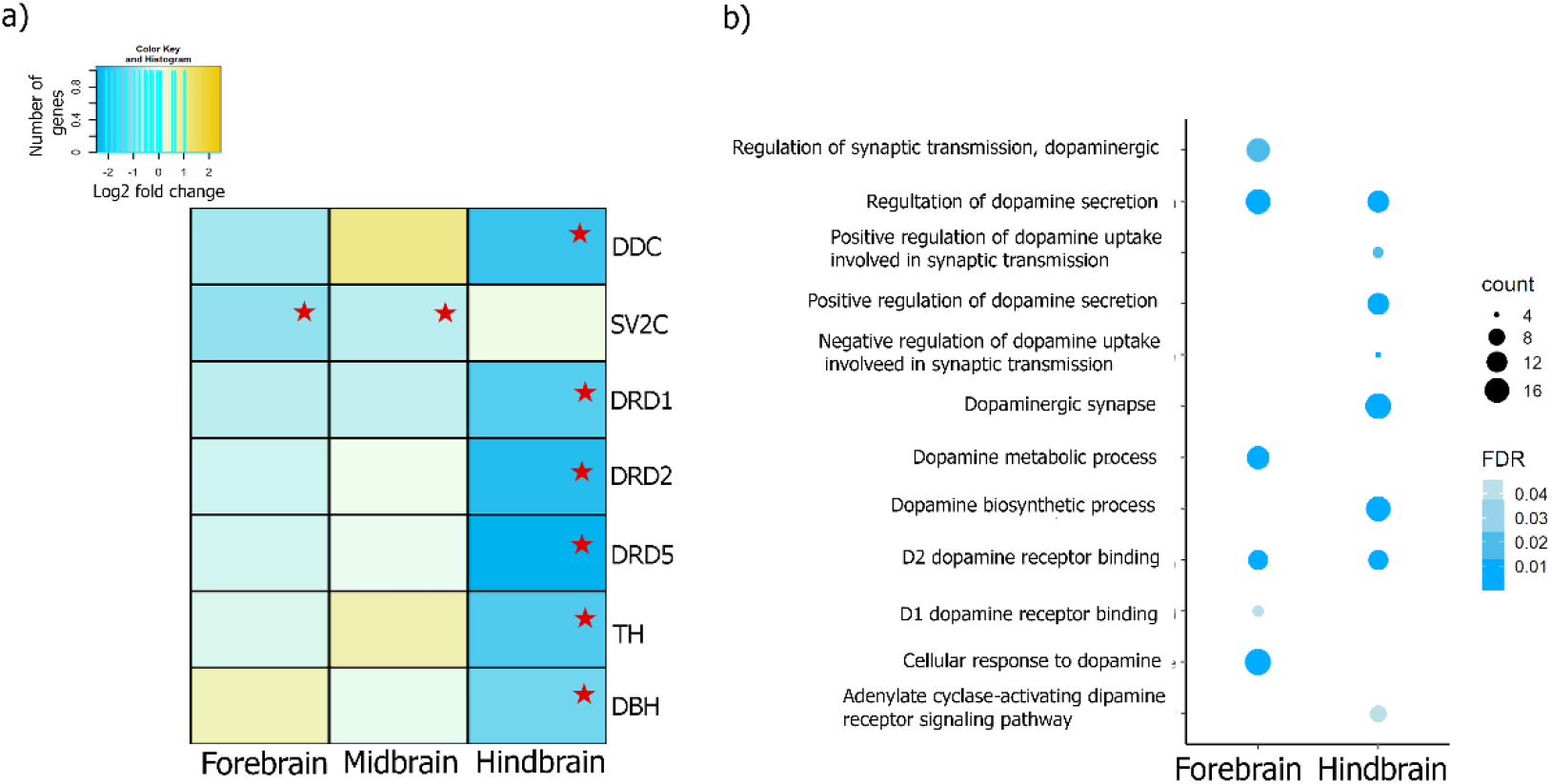
a) Dopaminergic pathway differential gene expression in the three brain regions of *L. dimidiatus* (genes obtained from KEGG PATHWAY Database-Dopaminergic synapse, entry: map04728). The colours presented correspond to log2fold changes. The red star represents significance. The legend indicates the reference values of log2fold changes for each DEG in the figure. b) Functional enrichment of Gene Ontology (GO) terms related to Dopamine activity in *L. dimidiatus* in the fore and hindbrain. No enrichment was found for the midbrain region. The size of the circles is proportional to the number of genes observed within each GO category, and the colour of the circles is proportional to the significance (FDR value).

Several differentially expressed genes (DEGs) were also found for *L. dimidiatus* related to the glutamatergic synapse pathway (figure 5; table S13 and figure S5). Many enrichments related to this pathway were exhibited in the brain of interacting *L. dimidiatus*, such as positive regulation of synaptic transmission glutamatergic, NMDA selective glutamate receptor signalling pathway, kainate selective glutamate receptor activity and AMPA glutamate receptor activity, among others with enrichments in the three regions of the brain (figure 5b). Interestingly, more functions were shared by the FB and HB, such as AMPA glutamate receptor complex, ionotropic glutamate receptor binding, NMDA glutamate receptor activity, glutamatergic neuron differentiation, glutamate binding and NMDA selective glutamate receptor complex (figure 5b). These functions stem from a total of 72 DEGs in the brain of *L. dimidiatus*, playing a role in the glutamatergic synapse processes (table S13). The differentially expressed genes within this pathway also followed our previous general pattern in which the HB region reports a higher number of significant DEGs (48), followed by the FB (46) and the MB (16; table S13). In particular, most of the ionotropic receptors (table S13were differentially expressed in the HB and FB except for GRIA3 and GRIA4, which were also differentially expressed in the MB. These receptors exhibited a downregulation pattern in all three regions of the brain when interacting with another species. In contrast, metabotropic receptors were differentially expressed in HB and FB during interaction condition (table S13); however, main receptors GRM1, 5 and 7 showed upregulation patterns in these two brain regions, while GRM3 and 4 were downregulated at HB.

**Figure 5.**
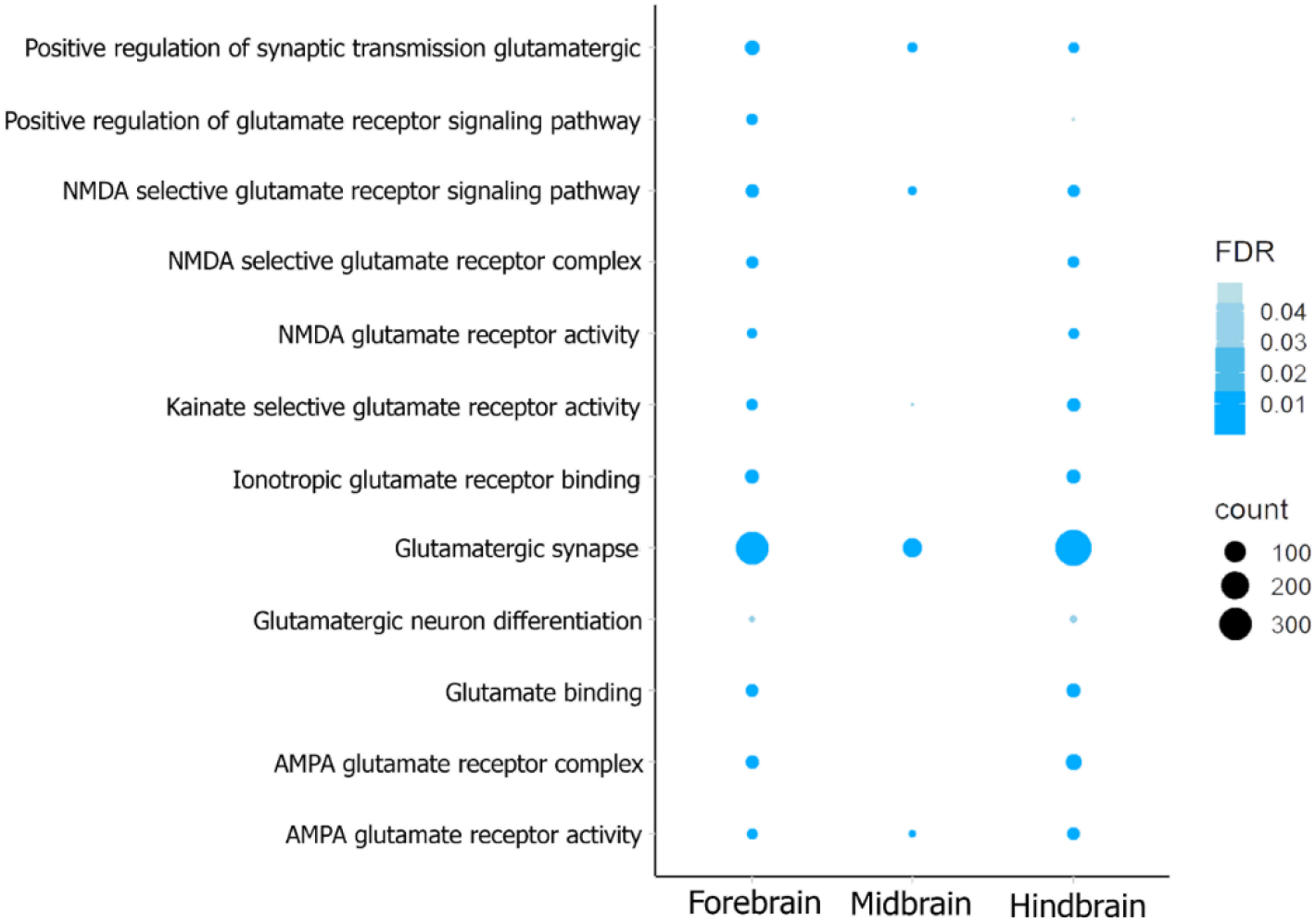
Functional enrichment of Gene Ontology (GO) terms related to the Glutamatergic synapse pathway in *L. dimidiatus* individuals in the fore, mid and hindbrain. The enrichments are based on differentially expressed genes of the comparison control vs interaction. The size of the circles is proportional to the number of genes observed within each GO category, and the colour of the circles is proportional to the significance (FDR value).

Only two DEGs related to the glutamate pathway were found in *A. leucosternon*: Glutamate carboxylase (DCE1) and Glutamate receptor ionotropic (GRID2). However, hormonal responses were enriched in the brain of *A. leucosternon* during the interaction condition (table S14). This was in concordance with the enrichment terms found in the FB region of this fish (Hormone activity, table S14). In particular, pituitary hormones such as Somatotropin (SOMA), Prolactin (PRL), Somatolactin (SOML2), Gonadotropins (GTHB1 and 2), Thyrotropin (TSHB) and Glycoproteins (GLHA) were differentially expressed and downregulated in the FB during the interaction condition, but highly expressed during the control (figure S6). Further differentially expressed genes were related with functions related of calcium-binding activity, oxygen transport and signalling were upregulated during the interaction in the FB (*i*.*e*. CHP2, FCRL5, PTPRF, ADA1B, MICU3, table S14).

## DISCUSSION

Distinct transcriptional and functional patterns were involved in the interaction behaviour of *L. dimidiatus* with its client fish *A. leucosternon*. This suggests that the social stimulus during the interaction has an effect on the transcription regulation of both fishes and that the larger number of altered molecular processes in the cleaner wrasse generates an adjustment of its behaviour. We observed a change in gene expression in functions related to behaviour in *L. dimidiatus* (e.g locomotory exploration behaviour) while interacting with the client. Genes underlying these functions included glutamatergic receptors (ionotropic and metabotropic), IEGs which are associated with learning processes and social plasticity (*i*.*e*. Proto-oncogene c-Fos (FOS) and cAMP response element-binding protein (CREB)) as well as dopaminergic pathway genes. The specific transcription found for *L. dimidiatus* suggests the activation of synaptic neurotransmission (*i*.*e*. glutamate) that can lead to mechanisms of memory consolidation in teleost fishes [39] and further learning and memory processes driven by dopamine and IEGs [40,41]. Previous studies have illustrated that the building blocks of cooperative behaviour in cleaner wrasses at ecological and physiological levels [22,24,42] include Arginine-vasotocin (AVT), Isotocin (IT) [43], Cortisol [22,44], Dopamine and Serotonin [23,45]. Hence, our results illustrate additional regulations at the transcriptional level of these social mechanisms when exposed to the presence of a new client. As for the client, the small number of genes and functions with altered expression found, suggests less transcriptional activity of genes regulating interaction behaviour in this species.

The underlying mechanisms in both species were exhibited in major areas of the brain, especially the Forebrain (FB) and Hindbrain (HB) revealing the extensive neural areas where behavioural modulation of cleaning interaction occurs. The HB region in teleost fishes (cerebellum, medulla oblongata and rhombomeres, [46]) is mainly known to be in charge of motor activity and autonomic responses (*i*.*e*. eye retraction, heart beating), but increasing evidence reveals that the cerebellum comprise major cognitive abilities of spatial learning and classical conditioning, such as the ability to learn from preceding events (stimulus) [47,48]. The large transcriptional changes exhibited for *L. dimidiatus* could be the result of a cascade of social stimulus processing underlined by glutamatergic neurotransmission receptors, IEGs and neurohormone receptors that modulate behaviour through Estrogen (ERR3) and Isotocin (ITR) receptors in the HB. Since most of the transcriptional regulation was found in this brain region, the molecular responses during the interaction may be generated on the basis of experience (*e*.*g*. recall previous interactions with clients [49,50]). *L. dimidiatus* is able to recognize clients and make decisions based on new social environments [20,22]. Therefore, the transcriptional regulation in the HB highlights mechanisms of learning and memory from possible previous events in this coral reef fish enabling new interactions.

Cognitive abilities such as memory and decision-making are developed in the FB region (or telencephalon, divided into diencephalon and secondary prosencephalon), which is well known for controlling motivated behaviour, memory, instinct modulation and decision-making [46]. During the interaction, we found that short and long-term memory consolidation processes as well as synaptic plasticity play an important role. In particular, IEGs such as Homer Scaffold Protein 1 (HOMER1) and neuronal PAS domain protein 4a (NPS4L) are involved in synapse formation and the expression of activity-dependent genes such as protein kinases. In addition, voltage-dependent calcium channel subunits (*i*.*e*. CAC1A) were differentially expressed in this brain region accompanied by several glutamatergic pathway genes (*e*.*g*. GRIK1, GRM5, NMDE1), suggesting an alteration of mechanisms of signalling that lead to changes in gene expression of excitatory neurons by synaptic activity [51,52]. This may be essential for the cleaner wrasse to consolidate relationships and memories with clients in the long-term. The expression of these genes has been previously observed in zebrafish during the establishment of social hierarchies between behavioural phenotypes (‘winners’ and ‘losers’ from confronting interactions; ‘mirror-fighters’; and non-interacting fish [5,53,54]), thus this shows that processes of memory making, and plasticity are essential also to the cleaning interaction behaviour of *L. dimidiatus*.

Additional mechanisms underlying cognitive functions were also identified through the differential expression of Progesterone and Estrogen receptors, which are known to regulate learning and memory processes in teleost fish when expressed in the hippocampus (FB) [42]. High mRNA levels of Estrogen receptor have been shown to modulate dominant behaviour in female zebrafish, causing changes in social status dynamics [55]. However, we found estrogen receptor gene expression to be downregulated during the interaction of the cleaner wrasse with its client, which could suggest a reduction in dominant behaviour or possibly a submissive approach when facing a client. On the other hand, the downregulation of the progesterone receptor in the FB suggests modulation of higher brain functions such as recognition, social relationships, learning and memory when interacting with a client. It has been demonstrated that the expression of Androgen, Progesterone and Estrogen receptors in *A. burtoni* lead to the regulation of complex social behaviour, behavioural plasticity, and the evaluation of rewarding stimulus between dominant and subordinate males [56,57]. Consequently, the differential expression of estrogen, progesterone and neurohormone receptors like Isotocin receptor (ITR) shown here in the HB and FB is a core behavioural mechanism regulating the cooperative behaviour of a highly social fish such as *L. dimidiatus* when interacting with client species.

To further understand how changes in gene expression mediate social cognition, IEGs can provide insights into rapid shifts in behavioural states [1]. In fact, *L. dimidiatus* differentially expressed IEGs in the brain during the interaction indicating an important activation of transcription factors and their participation in the transduction of signals during the interaction. For instance, the expression of *c-fos* (FOS) has been reported in the hippocampus region (FB) of cichlid fish *A. burtoni* during social and territorial interactions of males, indicating regulation of memory, spatial processing and social recognition when interacting with non-dominant males [58]. Thus, the upregulation of *c-fos* in the FB of *L. dimidiatus* may be suggesting similar neural mechanisms from this part of the brain, but in our case, they may be activated to recognize and interact with its clients. In addition, CREB regulatory factor (CREB) is critical for the consolidation of memory (long term potentiation or LTP, [51]) and was also found differentially expressed in the cleaner wrasse. Studies examining the molecular mechanisms of learning and memory in feeding and consolidation of new habits using mandarin fish (*Siniperca chuatsi*) also revealed differential expression of this gene when fish learn to eat dead prey after a training period [59]. Therefore, the differential expression of CREB in the brain of *L. dimidiatus* may suggest the activation of downstream processes of memory in the FB and MB when interacting with clients, which correspond to areas where LTP and associative learning occur in teleost fishes [46]. Consolidation of memory and learning is important for the cleaner wrasse as it can choose to clean, cheat or judge whether to provide tactile stimulation [45]. Here we observed specific molecular signatures of IEGs such as transcription factors *c-fos* and CREB, suggesting that social cognition, memory and learning processes are activated in the brain of this species during interaction with clients.

Dopamine is one of the major molecules related to signalling in the brain of vertebrates, and it is well-known to be involved in the modulation of animal behaviour and cognition [60]. We found the dopamine pathway is also one of the molecular regulators participating in the cooperative behaviour of *L. dimidiatus*. Dopamine signals are released upon specific stimuli [61], adjusting the way *L. dimidiatus* consolidates interactions with the clients as well as for associative learning [45,62]. For instance, disruption with antagonists of dopamine receptors D1 and D2 was shown to reduce levels of Dopamine transmission and increased the duration and frequency of cleaner wrasses tactile stimulation events during social encounters with clients [63]. In fact, D1 and D2 were downregulated in the HB, which emphasizes the role of dopamine in the modulation of behaviour during social interactions. Since animals need to re-evaluate decisions and make behavioural adjustments to new events (usually based on experience) [47,50,64], the downregulation of dopamine receptors in this region may be due to a response of dopamine signalling upon a new stimulus in *L. dimidiatus* (having another fish in the same tank). Consequently, the dopamine pathway expression patterns suggest that the cleaner wrasse may be increasing its willingness to interact with the client and possibly promote major tactile stimulation and negotiation. This result was also corroborated by our behavioural data since the 73.4% of the total cleaner-client interactions (N=327) were initiated only by the cleaner.

Lastly, the formation of cognitive abilities in the cleaner wrasse relied on the transcription of glutamate, which is the primary excitatory neurotransmitter in vertebrate brains [65]. Glutamatergic synapses are formed in specific sites of the brain where memory consolidation, synaptic plasticity, and storage occur [39]. We found glutamate-related genes differentially expressed in all three brain regions of *L. dimidiatus* during its interaction with the client, indicating a key role of this pathway in the mediation of synaptic transmission, activation of neurons and synapse plasticity. Differential expression of the two major groups of glutamatergic receptors (ionotropic and metabotropic), further suggests a contribution in the activation of long-term potentiation (LTP) processes and synaptic plasticity. In zebrafish, elevated expression of these receptors are associated with learnt behaviour to respond to electric shocks [66], while its distribution and differential expression defines neural connectivity and plasticity during development [67]. Further evidence comes from other fishes such as *Apteronotus leptorhynchus* in which the distribution of ionotropic receptors in the FB and HB suggest their contribution to learning and memory processes [68]. Consequently, our study suggests that LTP processes and plasticity may be occurring in the HB and FB regions in *L. dimidiatus* during the interaction and the glutamatergic synapse pathway plays an important role in external stimulus processing.

The molecular signatures underlying the interaction behaviour in the bluestreak cleaner wrasse were not evident in its client, the powder-blue surgeonfish. In the client we observed differential expression of several hormones in the FB such as Somatotropin releasing-hormone (SOMA), Prolactin (PRL), Somatolactin (SOML2), Pro-opiomelanocortin (POMC), among others, suggesting the transcriptional activity of the Hypothalamic-Pituitary-Thyroid (HPT) Axis [69]. While HPT in teleost fishes is mainly known for controlling development and growth through the secretion of these hormones [69–71], they also play a role in several behavioural aspects of fishes [72,73]. For instance, SOMA and PRL can regulate locomotion, feeding behaviour and cognitive functions in zebrafish [66,74] and aggression in rainbow trout [75]. SOML2 regulate stress in Atlantic cod and flounder [70] while POMC control levels of cortisol (CRH) leading to stress and food intake disruption in rainbow trout [76]. Consequently, as the HPT hormones found in our client fish had a significantly lower expression during the interaction, we can hypothesize a decrease in stress in *A. leucosternon* and therefore a change in the behaviour towards the cleaner wrasse and. In fact, *L. dimidiatus* is known to be able to provide stress relief to clients by lowering cortisol levels through physical contact (known as tactile stimulation) [16], and our behavioural trials support this since cleaners engaged on average in 3 ± 3.4 tactile stimulation events during the interactions. Consequently, since physical contact from *L. dimidiatus* can reduce stress in clients and a downregulation of HPT hormones related with stress was observed during the interaction, our results suggest stress-relief behaviour in the client fish.

In conclusion, differences in gene expression patterns in both fishes were noticeable, being *L. dimidiatus* the species with large transcriptional reprogramming and associated functions compared to *A. leucosternon*. During their interaction, immediate early genes were altered in expression in *L. dimidiatus* suggesting learning, memory processes and synaptic plasticity. Moreover, hormone activity including Estrogen, Progesterone, Isotocin receptors and Dopamine underlined important processes of associative learning, memory, decision-making and social plasticity mainly in the FB and HB. Finally, adjustments in the transcription of glutamatergic synapses suggested consolidation of long-term memory in the cleaner wrasse that may be key for longstanding cleaning relationships with clients. In contrast, in *A. leucosternon* the major molecular signal corresponded to a decreased transcription of Hypothalamic-Pituitary-Thyroid (HPT) hormone genes indicating stress reduction during the interaction. Our results highlight the functional genetics of cooperative behaviour in cleaner wrasses and the transcriptional changes in cognitive abilities implemented during the cleaning mutualism of *L. dimidiatus*.

## ACKNOWLEDGEMENTS

We would like to acknowledge Lígia Cascalheira and Dr Tiago Repolho for their help with the maintenance of aquatic systems throughout the experiment. We would like to thank Celia Schunter’s Lab members at HKU Sneha Suresh and Jingliang Kang, for their help building the bioinformatic pipeline and Natalia Petit, Arthur Chung and Jade Sourisse for engaging in stimulating discussions and comments to this work.

This work was supported by the Hong Kong Research Grant Committee Early Career Scheme fund 27107919 (CS) and CS’s HKU start-up fund (S.R. postgraduate studentship), the King Abdullah University of Science and Technology (TR & CS). Portuguese national funds funded this study through FCT–Fundação para a Ciência e Tecnologia, I.P., within the project PTDC/MAR-EST/5880/2014 (MUTUALCHANGE: Bio-ecological responses of marine cleaning mutualisms to climate change) to JRP and RR and the strategic project UID/MAR/04292/2020. JRP is currently supported by project ASCEND— PTDC/BIA-BMA/28609/2017 co-funded by FCT–Fundação para a Ciência e Tecnologia, I.P, Programa Operacional Regional de Lisboa, Portugal 2020 and the European Union Regional Development Fund within the project LISBOA-01-0145-FEDER-028609.

## DATA ACCESSIBILITY AND BENEFIT-SHARING STATEMENT

Raw sequencing files and *de novo* transcriptome assemblies (*L. dimidiatus* and *A. leucosternon*) have been submitted under the NCBI Bioproject number PRJNA726349. Reviewer link: https://dataview.ncbi.nlm.nih.gov/object/PRJNA726349?reviewer=lul18eg46dl6tjfnu5nhcaaigr. Benefits from this research accrue from the sharing of our data and results on public data-bases as described above.

## AUTHOR CONTRIBUTIONS

JRP built the experimental setup with input from RR. JRP & CS designed the project. JRP provided maintenance of aquarium setups, performed the behavioural assays and sampled the fish brains. EO and JRP analysed the behavioural videos and data. CS, with support from TR, conducted laboratory work and prepared samples for sequencing. SRC analysed data with input from CS. SRC and CS wrote the first draft, and all author edited and approved the final manuscript.

## REFERENCES

1. Oliveira RF. 2012 Social plasticity in fish: integrating mechanisms and function. J. Fish Biol. 81, 2127–2150. (doi:10.1111/j.1095-8649.2012.03477.x)

2. O’Connell LA, Hofmann HA. 2011 Genes, hormones, and circuits: An integrative approach to study the evolution of social behavior. Front. Neuroendocrinol. 32, 320–335. (doi:10.1016/j.yfrne.2010.12.004)

3. Oliveira RF. 2013 Mind the fish: zebrafish as a model in cognitive social neuroscience. Front. Neural Circuits 7, 1–15. (doi:10.3389/fncir.2013.00131)

4. Hofmann HA et al. 2014 An evolutionary framework for studying mechanisms of social behavior. Trends Ecol. Evol. 29, 581–589. (doi:10.1016/j.tree.2014.07.008)

5. Maruska K, Soares M, Lima-Maximino M, Henrique de Siqueira-Silva D, Maximino C. 2019 Social plasticity in the fish brain: Neuroscientific and ethological aspects. Brain Res. 1711, 156–172. (doi:10.1016/j.brainres.2019.01.026)

6. O’Connell LA, Hofmann HA. 2011 The Vertebrate mesolimbic reward system and social behavior network: A comparative synthesis. J. Comp. Neurol. 519, 3599–3639. (doi:10.1002/cne.22735)

7. Teles MC, Almeida O, Lopes JS, Oliveira RF. 2015 Social interactions elicit rapid shifts in functional connectivity in the social decision-making network of zebrafish. Proc. R. Soc. B Biol. Sci. 282. (doi:10.1098/rspb.2015.1099)

8. Rittschof CC et al. 2014 Neuromolecular responses to social challenge: Common mechanisms across mouse, stickleback fish, and honey bee. Proc. Natl. Acad. Sci. U. S. A. 111, 17929–17934. (doi:10.1073/pnas.1420369111)

9. Kasper C, Colombo M, Aubin-horth N, Taborsky B. 2018 Physiology & Behavior Brain activation patterns following a cooperation opportunity in a highly social cichlid fi sh. Physiol. Behav. 195, 37–47. (doi:10.1016/j.physbeh.2018.07.025)

10. Robinson GE, Fernald RD, Clayton DF. 2008 Genes and Social Behavior. Science (80-.). 322, 896–900. (doi:10.1126/science.1159277)

11. Barron AB, Robinson GE. 2008 The utility of behavioral models and modules in molecular analyses of social behavior. Genes, Brain Behav. 7, 257–265. (doi:10.1111/j.1601-183X.2007.00344.x)

12. Qiu Y-Q. 2013 KEGG Pathway Database. Encycl. Syst. Biol., 1068–1069. (doi:10.1007/978-1-4419-9863-7_472)

13. Bloch G, Grozinger CM. 2011 Social molecular pathways and the evolution of bee societies. Philos. Trans. R. Soc. B Biol. Sci. 366, 2155–2170. (doi:10.1098/rstb.2010.0346)

14. Waldie PA, Blomberg SP, Cheney KL, Goldizen AW, Grutter AS. 2011 Long-term effects of the cleaner fish Labroides dimidiatus on coral reef fish communities. PLoS One 6. (doi:10.1371/journal.pone.0021201)

15. Grutter AS. 1999 Cleaner fish really do clean. Nature (doi:10.1038/19443)

16. Soares M, Oliveira RF, Ros AFH, Grutter AS, Bshary R. 2011 Tactile stimulation lowers stress in fish. Nat. Commun. 2, 534–535. (doi:10.1038/ncomms1547)

17. Soares M, Gerlai R, Maximino C. 2018 The integration of sociality, monoamines and stress neuroendocrinology in fish models: applications in the neurosciences. J. Fish Biol. 93, 170–191. (doi:10.1111/jfb.13757)

18. Grutter A. 1996 Parasite removal rates by the cleaner wrasse Labroides dimidiatus. Mar. Ecol. Prog. Ser. 130, 61–70. (doi:10.3354/meps130061)

19. Grutter AS. 1997 Effect of the removal of cleaner fish on the abundance and species composition of reef fish. Oecologia 111, 137–143. (doi:10.1007/s004420050217)

20. Tebbich S, Bshary R, Grutter A. 2002 Cleaner fish Labroides dimidiatus recognise familiar clients. Anim. Cogn. 5, 139–145. (doi:10.1007/s10071-002-0141-z)

21. Pinto A, Oates J, Grutter A, Bshary R. 2011 Cleaner wrasses Labroides dimidiatus are more cooperative in the presence of an audience. Curr. Biol. 21, 1140–1144. (doi:10.1016/j.cub.2011.05.021)

22. Soares M. 2017 The Neurobiology of Mutualistic Behavior: The Cleanerfish Swims into the Spotlight. Front. Behav. Neurosci. 11, 1–12. (doi:10.3389/fnbeh.2017.00191)

23. Paula JR, Messias J, Grutter A, Bshary R, Soares M. 2015 The role of serotonin in the modulation of cooperative behavior. Behav. Ecol. 26, 1005–1012. (doi:10.1093/beheco/arv039)

24. de Abreu MS, Maximino C, Cardoso SC, Marques CI, Pimentel AFN, Mece E, Winberg S, Barcellos LJG, Soares MC. 2020 Dopamine and serotonin mediate the impact of stress on cleaner fish cooperative behavior. Horm. Behav. 125, 104813. (doi:10.1016/j.yhbeh.2020.104813)

25. Triki Z, Bshary R, Grutter AS, Ros AFH. 2017 The arginine-vasotocin and serotonergic systems affect interspecific social behaviour of client fish in marine cleaning mutualism. Physiol. Behav. 174, 136–143. (doi:10.1016/j.physbeh.2017.03.011)

26. Triki Z, Bshary R. 2021 Sex differences in the cognitive abilities of a sex-changing fish species Labroides dimidiatus. (doi:10.1098/rsos.210239)

27. Paula JR, Repolho T, Pegado MR, Thörnqvist P-O, Bispo R, Winberg S, Munday PL, Rosa R. 2019 Neurobiological and behavioural responses of cleaning mutualisms to ocean warming and acidification. Sci. Rep. 9, 1–10. (doi:10.1038/s41598-019-49086-0)

28. Soares M, Bshary R, Cardoso SC, Côté IM. 2008 The meaning of jolts by fish clients of cleaning gobies. Ethology 114, 209–214. (doi:10.1111/j.1439-0310.2007.01471.x)

29. Grutter AS. 2004 Cleaner fish use tactile dancing behavior as a preconflict management strategy. Curr. Biol. 14, 1080–1083. (doi:10.1016/j.cub.2004.05.048)

30. Friard O, Gamba M. 2016 BORIS: a free, versatile open-source event-logging software for video/audio coding and live observations. Methods Ecol. Evol. 7, 1325–1330. (doi:10.1111/2041-210X.12584)

31. Andrews S. 2010 Babraham Bioinformatics - FastQC: A Quality Control tool for High Throughput Sequence Data. See http://www.bioinformatics.babraham.ac.uk/projects/fastqc/ (accessed on 30 July 2020).

32. Bolger AM, Lohse M, Usadel B. 2014 Trimmomatic: A flexible trimmer for Illumina sequence data. Bioinformatics 30, 2114–2120. (doi:10.1093/bioinformatics/btu170)

33. Haas BJ et al. 2013 De novo transcript sequence reconstruction from RNA-seq using the Trinity platform for reference generation and analysis. Nat. Protoc. 8, 1494–1512. (doi:10.1038/nprot.2013.084)

34. Langmead B, Salzberg SL. 2012 Fast gapped-read alignment with Bowtie 2. Nat. Methods 9, 357–359. (doi:10.1038/nmeth.1923)

35. Waterhouse RM, Seppey M, Simao FA, Manni M, Ioannidis P, Klioutchnikov G, Kriventseva E V., Zdobnov EM. 2018 BUSCO applications from quality assessments to gene prediction and phylogenomics. Mol. Biol. Evol. 35, 543–548. (doi:10.1093/molbev/msx319)

36. Götz S et al. 2008 High-throughput functional annotation and data mining with the Blast2GO suite. Nucleic Acids Res. 36, 3420–3435. (doi:10.1093/nar/gkn176)

37. Love MI, Huber W, Anders S. 2014 Moderated estimation of fold change and dispersion for RNA-seq data with DESeq2. Genome Biol. 15, 550. (doi:10.1186/s13059-014-0550-8)

38. Schunter C, Jarrold MD, Munday PL, Ravasi T. 2021 Diel CO2 fluctuations alter the molecular response of coral reef fishes to ocean acidification conditions. bioRxiv (doi:10.1101/2021.05.17.444406)

39. Bajaffer A, Mineta K, Gojobori T. 2021 Evolution of memory system-related genes. FEBS Open Bio 11, 3201–3210. (doi:10.1002/2211-5463.13224)

40. O’Connell LA, Fontenot MR, Hofmann HA. 2011 Characterization of the dopaminergic system in the brain of an African cichlid fish, Astatotilapia burtoni. J. Comp. Neurol. 519, 75–92. (doi:10.1002/cne.22506)

41. Oliveira RF. 2012 Social plasticity in fish : integrating mechanisms. J. Fish Biol. 81, 2127–2150. (doi:10.1111/j.1095-8649.2012.03477.x)

42. Soares M, Bshary R, Fusani L, Goymann W, Hau M, Hirschenhauser K, Oliveira RF. 2010 Hormonal mechanisms of cooperative behaviour. Philos. Trans. R. Soc. B Biol. Sci. 365, 2737–2750. (doi:10.1098/rstb.2010.0151)

43. Soares M, Bshary R, Mendonça R, Grutter AS, Oliveira RF. 2012 Arginine vasotocin regulation of interspecific cooperative behaviour in a cleaner fish. PLoS One 7, 39583. (doi:10.1371/journal.pone.0039583)

44. Soares M, Cardoso SC, Grutter AS, Oliveira RF, Bshary R. 2014 Cortisol mediates cleaner wrasse switch from cooperation to cheating and tactical deception. Horm. Behav. 66, 346–350. (doi:10.1016/j.yhbeh.2014.06.010)

45. Messias J, Santos TP, Pinto M, Soares MC. 2016 Stimulation of dopamine D1 receptor improves learning capacity in cooperating cleaner fish. Proc. R. Soc. B Biol. Sci. 283. (doi:10.1098/rspb.2015.2272)

46. Vernier P. 2016 The Brains of Teleost Fishes. (doi:10.1016/B978-0-12-804042-3.00004-X)

47. Terry WS. 2021 Classical Conditioning. In Learning and Memory, pp. 76–112. (doi:10.4324/9781315665023-10)

48. Rodríguez F, Broglio C, Durán E, Gómez A, Salas C. 2007 Neural Mechanisms of Learning in Teleost Fish. Fish Cogn. Behav., 243–277. (doi:10.1002/9780470996058.ch13)

49. Brown C, Laland K, Krause J, Blbk FM, June B, Count C. 2011 Fish Cognition and Behavior. (doi:10.1016/j.scitotenv.2015.08.111)

50. Odling-Smee L, Braithwaite VA. 2003 The role of learning in orientation. Fish Fish. 4, 235–246.

51. Alberini CM. 2009 Transcription factors in long-term memory and synaptic plasticity. Physiol. Rev. 89, 121–145. (doi:10.1152/physrev.00017.2008)

52. Limbäck-Stokin K, Korzus E, Nagaoka-Yasuda R, Mayford M. 2004 Nuclear calcium/calmodulin regulates memory consolidation. J. Neurosci. 24, 10858–10867. (doi:10.1523/JNEUROSCI.1022-04.2004)

53. Klaric T, Lardelli M, Key B, Koblar S, Lewis M. 2014 Activity-dependent expression of neuronal PAS domain-containing protein 4 (npas4a) in the developing zebrafish brain. Front. Neuroanat. 8, 1–13. (doi:10.3389/fnana.2014.00148)

54. Teles MC, Cardoso SD, Oliveira RF. 2016 Social Plasticity Relies on Different Neuroplasticity Mechanisms across the Brain Social Decision-Making Network in Zebrafish. Front. Behav. Neurosci. 10, 16. (doi:10.3389/fnbeh.2016.00016)

55. Filby AL, Paull GC, Bartlett EJ, Van Look KJW, Tyler CR. 2010 Physiological and health consequences of social status in zebrafish (Danio rerio). Physiol. Behav. 101, 576–587. (doi:10.1016/j.physbeh.2010.09.004)

56. Munchrath LA, Hofmann HA. 2010 Distribution of sex steroid hormone receptors in the brain of an african cichlid fish, Astatotilapia burtoni. J. Comp. Neurol. 518, 3302–3326. (doi:10.1002/cne.22401)

57. O’Connell LA, Ding JH, Hofmann HA. 2013 Sex differences and similarities in the neuroendocrine regulation of social behavior in an African cichlid fish. Horm. Behav. 64, 468–476. (doi:10.1016/j.yhbeh.2013.07.003)

58. Weitekamp CA, Hofmann HA. 2017 Neuromolecular correlates of cooperation and conflict during territory defense in a cichlid fish. Horm. Behav. 89, 145–156. (doi:10.1016/j.yhbeh.2017.01.001)

59. Dou Y, He S, Liang XF, Cai W, Wang J, Shi L, Li J. 2018 Memory function in feeding habit transformation of mandarin fish (Siniperca chuatsi). Int. J. Mol. Sci. 19. (doi:10.3390/ijms19041254)

60. Arias-Carrión O, Stamelou M, Murillo-Rodríguez E, Menéndez-Gonzalez M, Pöppel E. 2010 Dopaminergic reward system: A short integrative review. Int. Arch. Med. 3. (doi:10.1186/1755-7682-3-24)

61. Schultz W. 2006 Behavioral theories and the neurophysiology of reward. Annu. Rev. Psychol. 57, 87–115. (doi:10.1146/annurev.psych.56.091103.070229)

62. Dewitt EEJ. 2014 Neuroeconomics: A formal test of dopamine’s role in reinforcement learning. Curr. Biol. 24, R321–R324. (doi:10.1016/j.cub.2014.02.055)

63. Messias J, Paula JR, Grutter AS, Bshary R, Soares MC. 2016 Dopamine disruption increases negotiation for cooperative interactions in a fish. Sci. Rep. 6, 2–9. (doi:10.1038/srep20817)

64. Steinberg E, Keiflin R, Boivin J, Witten I, k D, Janak P. 2014 A Causal Link Between Prediction Errors, Dopamine Neurons and Learning. Nat. Neurosci. 16, 1–19. (doi:10.1038/nn.3413.A)

65. Meldrum BS. 2000 Glutamate as a neurotransmitter in the brain: review of physiology and pathology. J. Nutr. 130, 1007S–15S. (doi:10.1093/jn/130.4.1007S)

66. Studzinski ALM, Barros DM, Marins LF. 2015 Growth hormone (GH) increases cognition and expression of ionotropic glutamate receptors (AMPA and NMDA) in transgenic zebrafish (Danio rerio). Behav. Brain Res. 294, 36–42. (doi:10.1016/j.bbr.2015.07.054)

67. Hoppmann V, Wu JJ, Søviknes AM, Helvik JV, Becker TS. 2008 Expression of the eight AMPA receptor subunit genes in the developing central nervous system and sensory organs of zebrafish. Dev. Dyn. 237, 788–799. (doi:10.1002/dvdy.21447)

68. Weld MM, Kar S, Maler L, Quirion R. 1991 The distribution of Excitatory Amino Acid Binding Sites in the Brain of an Electric Fish, Apteronotus leptorhynchus. J. Chem. Neuroanat. 4, 39–61. (doi:10.1016/0891-0618(94)90024-8)

69. Blanton ML, Specker JL. 2007 The hypothalamic-pituitary-thyroid (HPT) axis in fish and its role in fish development and reproduction. Crit. Rev. Toxicol. 37, 97–115 (doi:10.1080/10408440601123529)

70. Kawauchi H, Sower SA, Moriyama S. 2009 Chapter 5. The Neuroendocrine Regulation of Prolactin and Somatolactin Secretion in Fish. In Fish Physiology, pp. 197–234. Elsevier Inc. (doi:10.1016/S1546-5098(09)28005-8)

71. Helmreich DL, Parfitt DB, Lu XY, Akil H, Watson SJ. 2005 Relation between the Hypothalamic-Pituitary-Thyroid (HPT) axis and the Hypothalamic-Pituitary-Adrenal (HPA) axis during repeated stress. Neuroendocrinology 81, 183–192. (doi:10.1159/000087001)

72. Jönsson E, Björnsson B. 2002 Physiological functions of growth hormone in fish with special reference to its influence on behaviour. Fish. Sci. 68, 742–748. (doi:10.2331/fishsci.68.sup1_742)

73. Zoeller RT, Tan SW, Tyl RW. 2007 General background on the hypothalamic-pituitary-thyroid (HPT) axis. Crit. Rev. Toxicol. 37, 11–53. (doi:10.1080/10408440601123446)

74. Björnsson B, Johansson V, Benedet S, Einarsdottir IE, Hildahl J, Agustsson T, Jönsson E. 2002 Growth hormone endocrinology of salmonids: Regulatory mechanisms and mode of action. Fish Physiol. Biochem. 27, 227–242. (doi:10.1023/B:FISH.0000032728.91152.10)

75. Trainor BC, Hofmann HA. 2006 Somatostatin Regulates aggressive behavior in an african cichlid fish. Endocrinology 147, 5119–5125. (doi:10.1210/en.2006-0511)

76. Doyon C, Gilmour KM, Trudeau VL, Moon TW. 2003 Corticotropin-releasing factor and neuropeptide Y mRNA levels are elevated in the preoptic area of socially subordinate rainbow trout. Gen. Comp. Endocrinol. 133, 260–271. (doi:10.1016/S0016-6480(03)00195-3)

