## Supplementary figures for "Neuro-molecular characterization of fish cleaning interactions"

**SUPPLEMENTARY MATERIAL**

**FIGURES:**

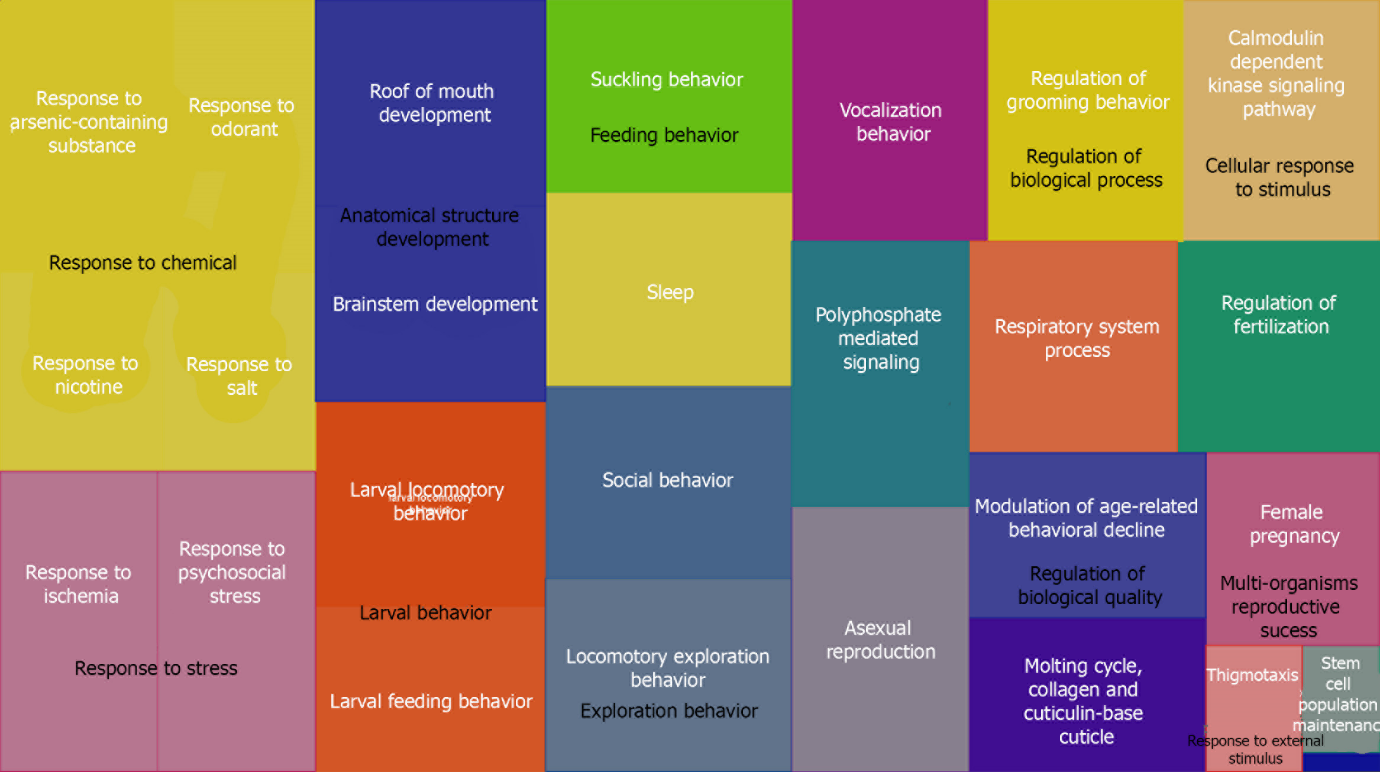

**Figure S1**. Gene Ontology treemap for *L. dimidiatus* representing the commonly significantly enriched functions in the forebrain region during the interaction treatment. Boxes with the same colour correspond to the upper-hierarchy GO-term and its title is found in the middle of each box.

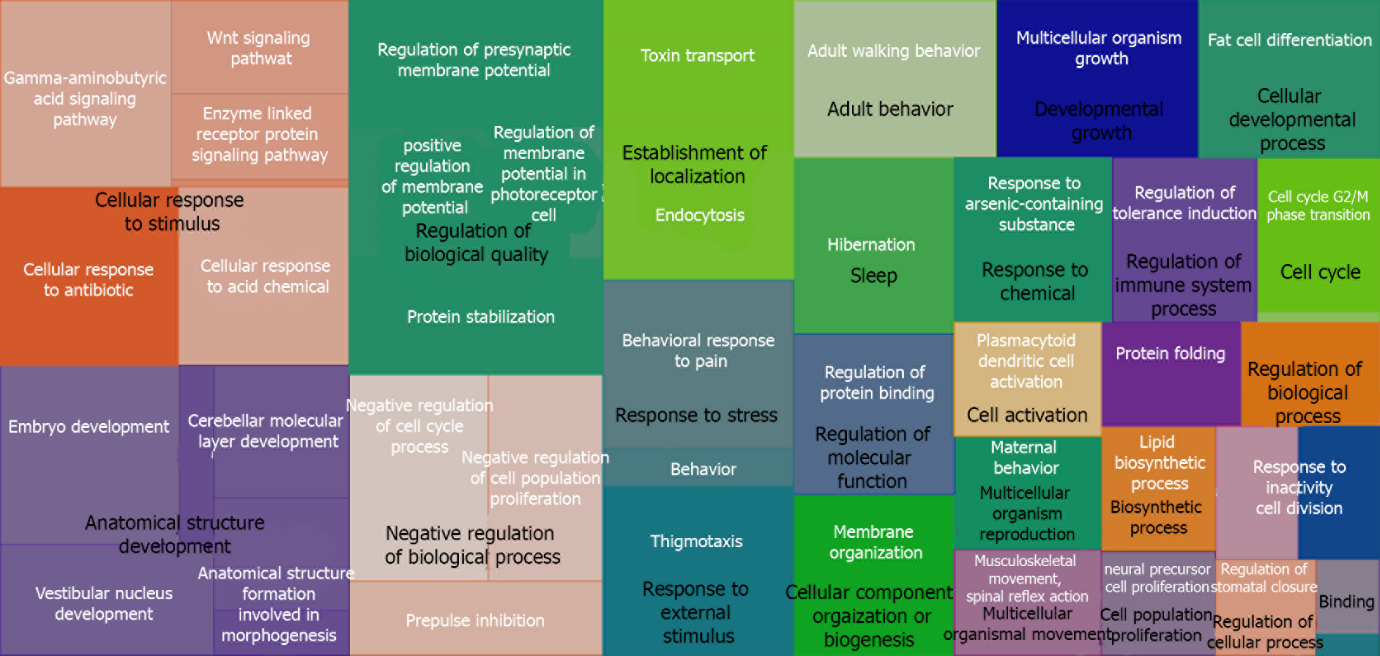

**Figure S2**. Gene Ontology treemap for *L. dimidiatus* representing the commonly significant enriched functions in the midbrain region during the interaction treatment. Boxes with the same colour correspond to the upper-hierarchy GO-term and its title is found in the middle of each box.

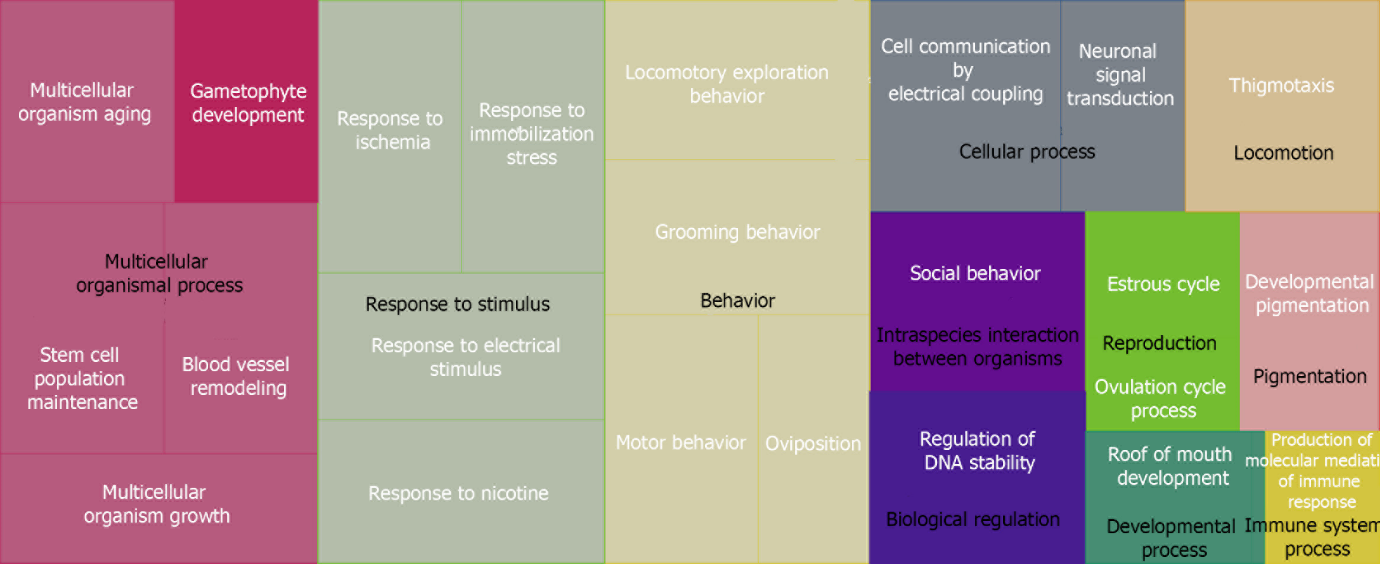

**Figure S3.** Gene Ontology treemap for *L. dimidiatus* representing the commonly significant enriched functions in the hindbrain region during the interaction treatment. Boxes with the same colour correspond to the upper-hierarchy GO-term and its title is found in the middle of each box.

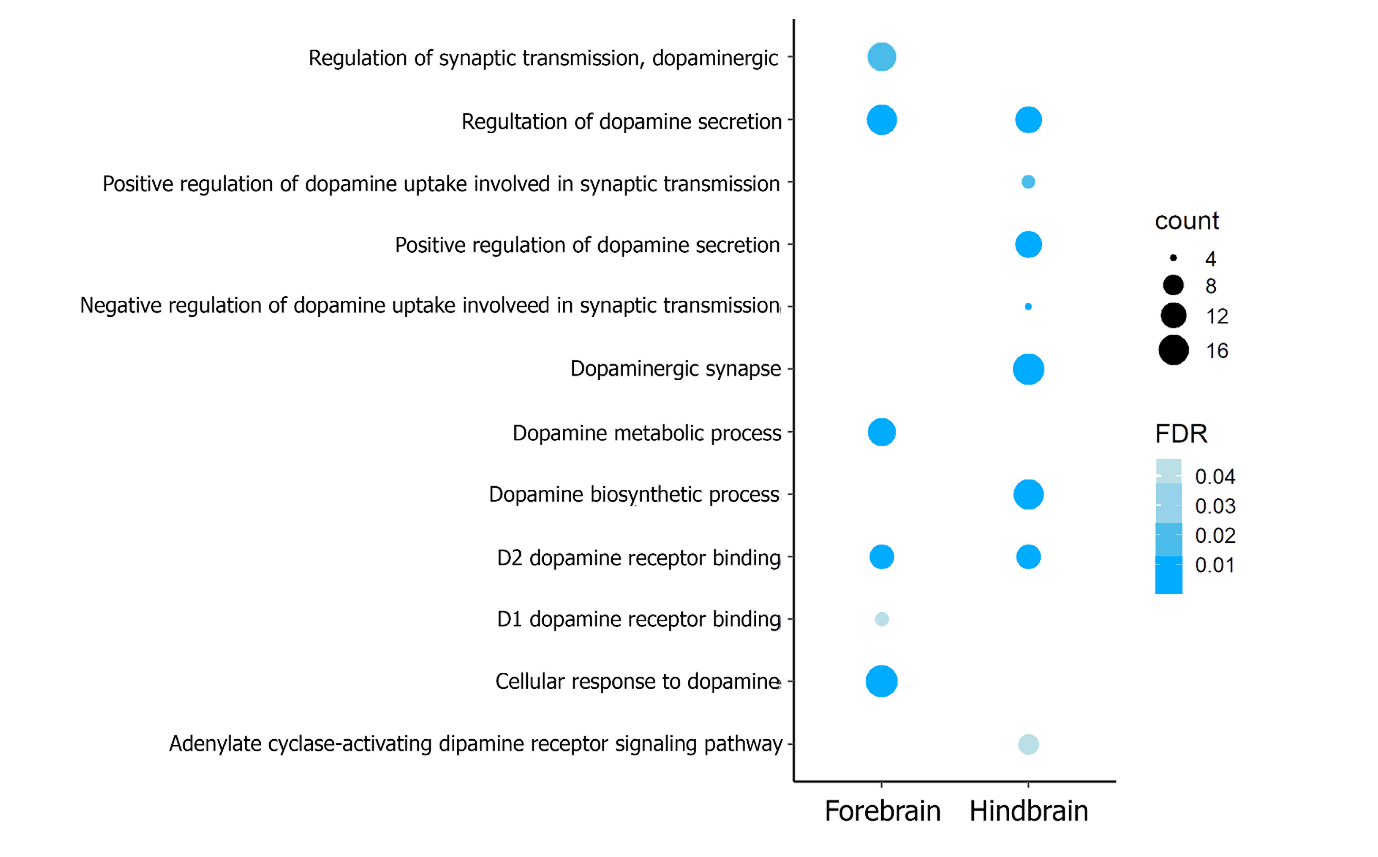

**Figure S4.** Functional enrichment of Gene Ontology (GO) terms related to Dopamine activity in *L. dimidiatus* in the fore and hindbrain. No enrichment was found for the midbrain region. The size of the circles is proportional to the number of genes observed within each GO category, and the colour of the circles is proportional to the significance (FDR value)

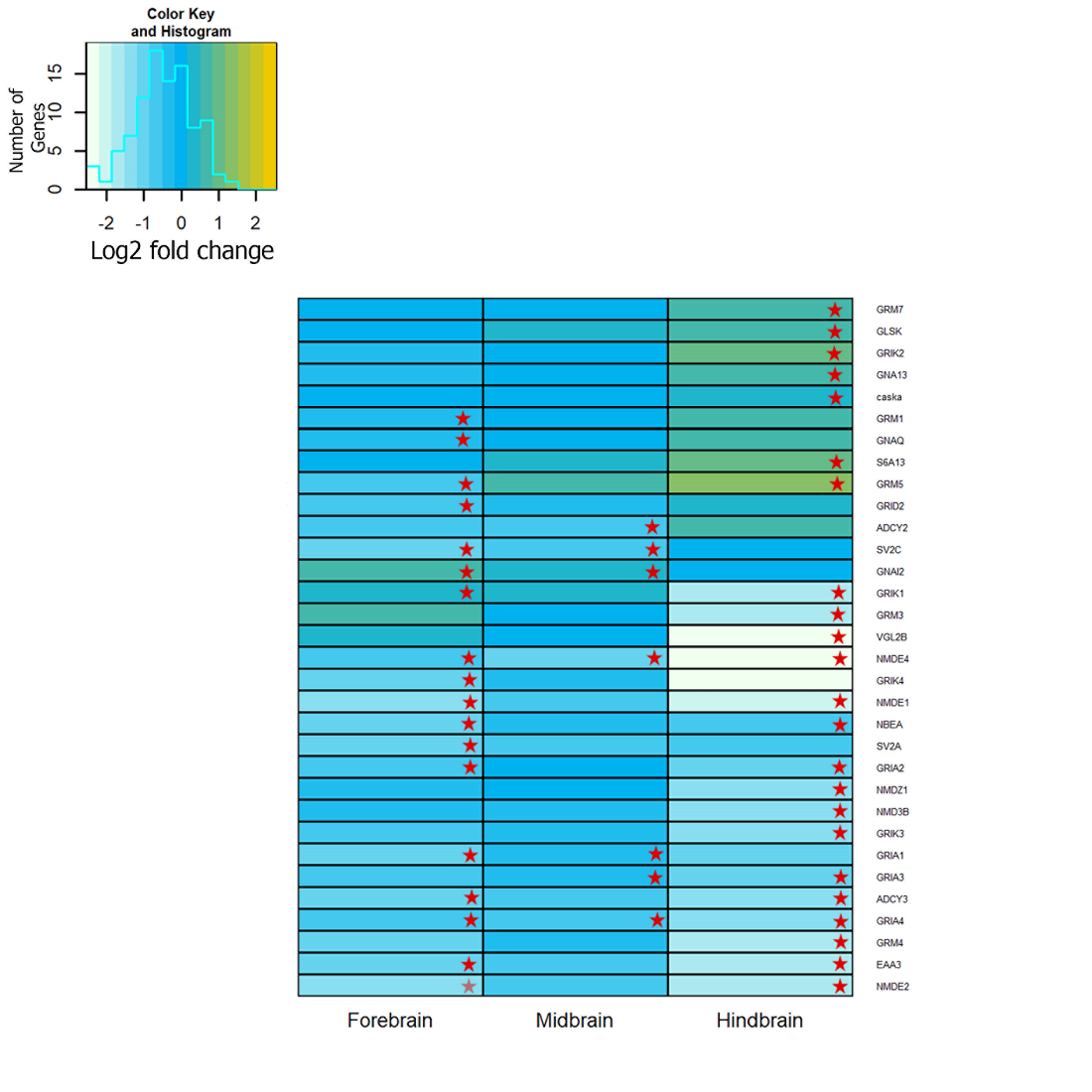

**Figure S5**. Comparative differential gene expression patterns of Glutamatergic synapse genes in the regions of the brain of *L. dimidiatus*. Red stars represent significance (padj<0.05), while colours represent log2fold change estimates (threshold of 0.3). Gene ADCY2 (Adenylate cyclase type 2), ADCY3 (Adenylate cyclase type 3), CA2D1 (Voltage-dependent calcium channel subunit alpha-2/delta-1), CA2D2 (Voltage-dependent calcium channel subunit alpha-2/delta-2), CABP7 (Calcium-binding protein 7), CAC1A (Voltage-dependent P/Q-type calcium channel subunit alpha-1A), CAC1B (Probable voltage-dependent N-type calcium channel subunit alpha-1B), CAC1C (Voltage-dependent L-type calcium channel subunit alpha-1C), CAC1D (Voltage-dependent L-type calcium channel subunit alpha-1D), CAC1G (Voltage-dependent T-type calcium channel subunit alpha-1G), CAC1H (Voltage-dependent T-type calcium channel subunit alpha-1H), CAC1I (Voltage-dependent T-type calcium channel subunit alpha-1I), CACB2 (Voltage-dependent L-type calcium channel subunit beta-2), CASKA (calcium/calmodulin dependent serine protein kinase), CCG2 (Voltage-dependent calcium channel gamma-2 subunit), CCG4	(Voltage-dependent calcium channel gamma-4 subunit), CCG5 (Voltage-dependent calcium channel gamma-5 subunit), CCG8 (Voltage-dependent calcium channel gamma-8 subunit) CPLX2 (Calcium/calmodulin-dependent protein kinase type 1), CSKP (Calcium/calmodulin-dependent protein kinase type 1D), EAA3 (Excitatory amino acid transporter 3), GABR1 (Gamma-aminobutyric acid type B receptor subunit 1), GABR2 (Gamma-aminobutyric acid type B receptor subunit 2), GBLP (Guanine nucleotide-binding protein subunit beta-2-like 1), GBRB4 (Gamma-aminobutyric acid receptor subunit beta-4), GBRG1 (Gamma-aminobutyric acid receptor subunit gamma-1), GBRP (Gamma-aminobutyric acid receptor subunit pi), GBRR1 (Gamma-aminobutyric acid receptor subunit rho-1), GBRR2 (Gamma-aminobutyric acid receptor subunit rho-2), GCR (Glucocorticoid receptor), GCYA1 (Guanylate cyclase soluble subunit alpha-1), GCYA2 (Guanylate cyclase soluble subunit alpha-2), GCYB1 (Guanylate cyclase soluble subunit beta-1), GLRA1 (Glycine receptor subunit alphaZ1), GLRA2 (Glycine receptor subunit alpha-2), GLRA4 (Glycine receptor subunit alpha-4), GLRB (Glycine receptor subunit beta), GLSK	(Glutaminase kidney isoform, mitochondrial), GNA13 (Guanine nucleotide-binding protein subunit alpha-13), GNAI1 (Guanine nucleotide-binding protein G(i) subunit alpha-1), GNAI2 (Guanine nucleotide-binding protein G(i) subunit alpha-2), GNAQ (BELL-associated factor 1), GNB5A (Guanine nucleotide-binding protein subunit beta-5a), GNL1	(Guanine nucleotide-binding protein-like 1), GRIA1 (Glutamate receptor 1), GRIA2 (Glutamate receptor 2), GRIA3 (Glutamate receptor 3), GRIA4 (Glutamate receptor 4), GRID2 (Glutamate receptor ionotropic, delta-2), GRIK1 (Glutamate receptor ionotropic, kainate 1, GRIK2 (Glutamate receptor ionotropic, kainate 2), GRIK3 (Glutamate receptor ionotropic, kainate 3), GRIK4 (Glutamate receptor ionotropic, kainate 4), GRM1 (Metabotropic glutamate receptor 1), GRM3 (Metabotropic glutamate receptor 3), GRM4 (Metabotropic glutamate receptor 4), GRM5 (Metabotropic glutamate receptor 5, GRM7 (Metabotropic glutamate receptor 7, KAPCA (cAMP-dependent protein kinase catalytic subunit alpha), KC2D2 (Calcium/calmodulin-dependent protein kinase type II delta 2 chain), KCC1D (Calcium/calmodulin-dependent protein kinase type 1D), KCC1G (Calcium/calmodulin-dependent protein kinase type 1G), KCC2A (Gamma-aminobutyric acid receptor subunit beta-3), NAC1	(Voltage-dependent P/Q-type calcium channel subunit alpha-1A), NBEA (Glucocorticoid receptor), NMD3B (Glutamate receptor ionotropic, NMDA 3B), NMDE1 (Glutamate receptor ionotropic, NMDA 2A), NMDE2 (Glutamate receptor ionotropic, NMDA 2B), NMDE4 (Glutamate receptor ionotropic, NMDA 2D), NMDZ1 (Glutamate receptor ionotropic, NMDA 1), S6A13 (Sodium- and chloride-dependent GABA transporter 2), SV2A (Synaptic vesicle glycoprotein 2A), SV2C (Synaptic vesicle glycoprotein 2C), VGL2B (Vesicular glutamate transporter 2.2). The legend indicates the reference values of log2fold changes for each DEG in the figure.

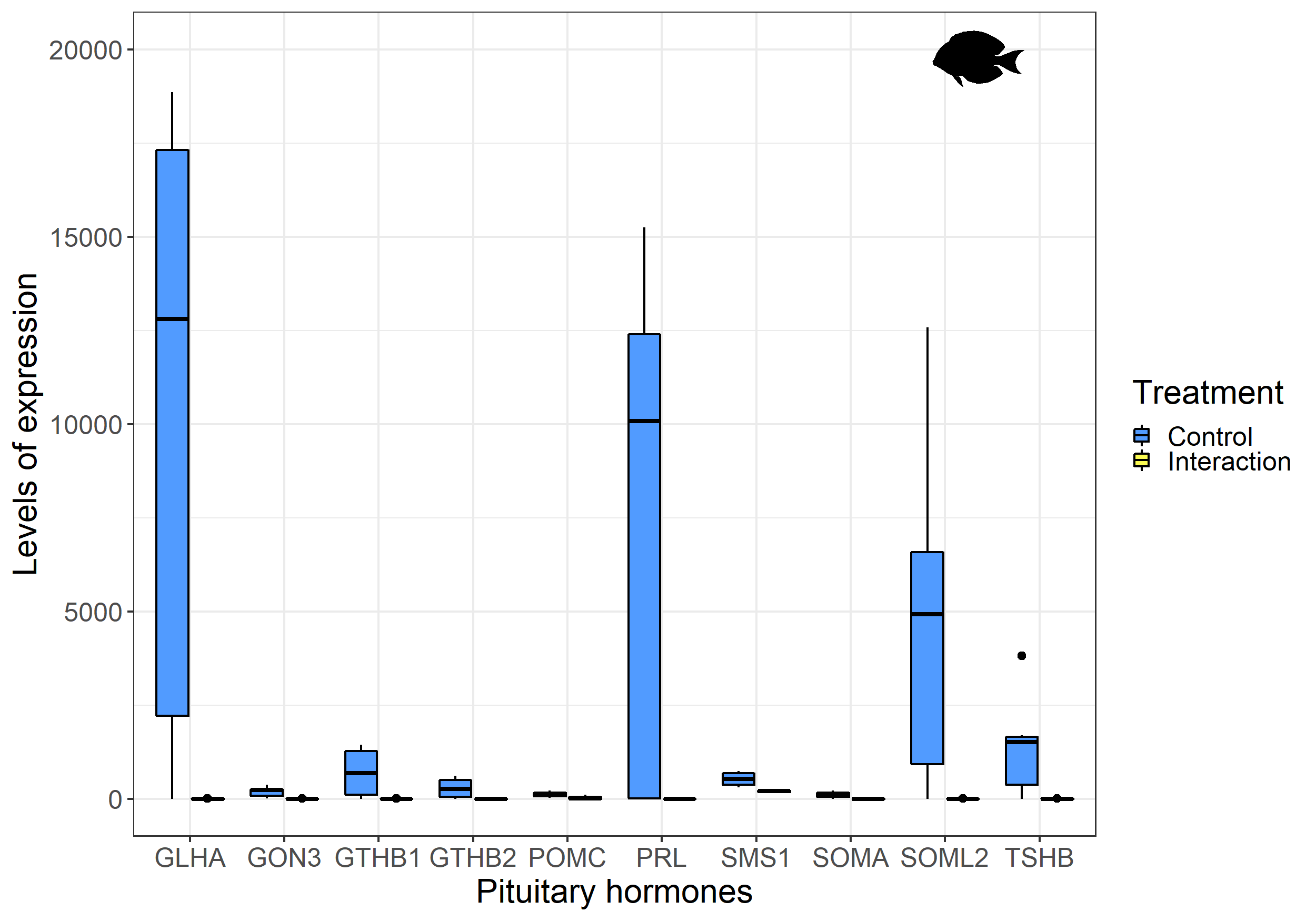

**Figure S6.** Gene expression levels of significant Hypothalamic-Pituitary-Thyroid (HPT) hormone genes in the forebrain region of *Acanthurus leucosternon* during the interaction with *L. dimidiatus*. Gene GLHA (Glycoprotein hormones alpha chain), GON3 (Progonadoliberin-3), GTHB1 (Gonadotropin subunit beta-1), GTHB2	(Gonadotropin subunit beta-2), POMC (Pro-opiomelanocortin), PRL (Prolactin), SMS1 (Somatostatin-1), SOMA (Somatotropin), SOML2 (Somatolactin-2), TSHB (Thyrotropin subunit beta).
